# Paternally-induced environmental programming of placenta development, offspring birth weight and health phenotypes in a mouse model

**DOI:** 10.1101/2023.12.07.570618

**Authors:** Elaine Chen, Raquel Santana da Cruz, Aallya Nascimento, Meghali Joshi, Duane Gischewski Pereira, Odalys Dominguez, Gabriela Fernandes, Megan Smith, Sara P.C. Paiva, Sonia de Assis

**Affiliations:** Department of Oncology, Lombardi Comprehensive Cancer Center, Georgetown University, Washington, DC, USA; Department of Obstetrics and Gynecology, Federal University of Minas Gerais (UFMG), Belo Horizonte, MG, Brazil

## Abstract

Mounting evidence suggests that environmentally induced epigenetic inheritance occurs in mammals and that traits in the progeny can be shaped by parental environmental experiences. Epidemiological studies link parental exposure to environmental toxicants, such as the pesticide DDT, to health phenotypes in the progeny, including low birth and increased risk of chronic diseases later in life. Here, we show that the progeny of male mice exposed to DDT in the pre-conception period are born smaller and exhibit sexual dimorphism in metabolic function, with male offspring developing both glucose intolerance and insulin resistance compared to controls. These phenotypes in DDT offspring were linked to reduced fetal growth and placenta size as well as placenta-specific reduction of glycogen levels and the nutrient sensor and epigenetic regulator OGT, with more pronounced phenotypes observed in male placentas. However, placenta-specific genetic reduction of OGT only partially replicates the metabolic phenotype observed in offspring of DDT-exposed males. Our findings reveal a role for paternal pre-conception environmental experiences in shaping placenta development and in fetal growth restriction. While many questions remain, our data raise the tantalizing possibility that placenta programming could be a mediator of environmentally induced intergenerational epigenetic inheritance of phenotypes and needs to be further evaluated.

## Introduction

In eutherian mammals such as human and rodents, the placenta connects the fetus to the mother and provides the nutrient and gas exchanges necessary for optimal fetal development ^1,2^. It also offers protection from the environment and endocrine support to the pregnancy ^1,2 3^. This extraembryonic organ is derived from the trophectoderm, the structure forming the outer cell layer in pre-implantation embryos at the blastocyst stage ^2^. The trophectoderm-derived cell lineage is the first lineage to appear during mammalian development. This cell lineage then gives rise to different placental trophoblast cell types. Trophoblast giant cells, for instance, are crucial for embryo implantation as they allow for infiltration into the maternal uterine epithelium and embryo attachment to the endometrium ^2^. Upon implantation, cells in the uterus lining undergo a decidualization process that is critical for normal placentation and embryonic growth and survival^4^.

Although there are organizational differences between the human and the mouse placenta, the role of the major placental structures is quite similar between these two species. They also share comparable gene and protein expression patterns and, consequently, mouse models are considered excellent surrogates to study many aspects of the human placenta^3^. In mice, the placenta contains three main layers. The decidua is derived from the maternal endometrium while the junctional and labyrinth zones are derived from the embryonic trophectoderm ^2^. Within the junctional zone, glycogen cells provide energetic and hormonal support for fetal growth. The labyrinth zone is the site of nutrient and gas exchange between mother and fetus ^2,3^. Defects in both the maternally and embryonically derived layers can cause fetal growth restriction ^2^. Abnormal placenta function not only impacts fetal growth but also offspring’s health after birth and throughout the lifespan ^3^. A link between placenta dysfunction and maternal health in pregnancy is also well-documented ^3,5^.

There is accumulating evidence for a placental role in epigenetic inheritance and intergenerational transmission of disease predisposition ^3^. Because of the close relationship between mother and fetus, many studies have documented the effects of maternal experiences in pregnancy on placenta development and function ^5 6^. Yet, despite the growing number of studies showing the importance of pre-conception paternal environmental exposures on offspring’s health, little is known about the impact of paternal experiences on placenta formation, although evidence is emerging ^7,8,9^. Given its embryonic origin, is reasonable to expect that both paternal and maternal factors play a part in placenta development. Surprisingly, seminal studies on imprinting suggest that the process of placenta development is primarily driven by the paternal genome and epigenome while maternal genes are more important in cell and tissue development of the embryo itself ^10,11^. In support of that, it has been reported that the paternal epigenome has a major impact in placenta formation and that paternally expressed genes are predominant in placenta ^12^.

Epidemiological studies link pre-conception paternal exposure to persistence organic pollutants and other environmental chemicals to low birth size in offspring in humans ^13,14 15 16^. Low birth weight is often a result of placenta defects, yet there is a lack of studies investigating the link between paternal exposure to environmental toxicants and placenta development.

Here, we developed a mouse model of preconception DDT exposure to investigate whether paternal environmental experiences reprogram placenta development, birth weight and later life health. We found that preconception paternal DDT exposure is linked to a decrease in birth weight and in placenta size as well as placenta-specific suppression of the nutrient sensor and epigenetic regulator OGT. Offspring of DDT-exposed fathers also show a sex-specific increase in metabolic dysfunction. However, placenta-specific genetic reduction of OGT only partially replicates the metabolic phenotype observed in offspring of DDT-exposed males.

## Results

### Paternal DDT exposure reduces offspring’s birth weight

When mated with unexposed female mice (**Fig. 1a**), DDT-exposed males produced offspring that were smaller at birth compared to controls (**Fig. 1b-c**). This is in agreement with human studies showing that parental DDT and other pesticides exposure is linked to low birth weight in offspring^13^. No significant differences in litter size or sex distribution were observed (**Fig. 1d-e**). As they aged, DDT offspring underwent a catch-up growth period and beginning at 3 weeks of age, no differences in body weight were observed between the groups (**Fig. 1f-g**).

**Figure 1.**
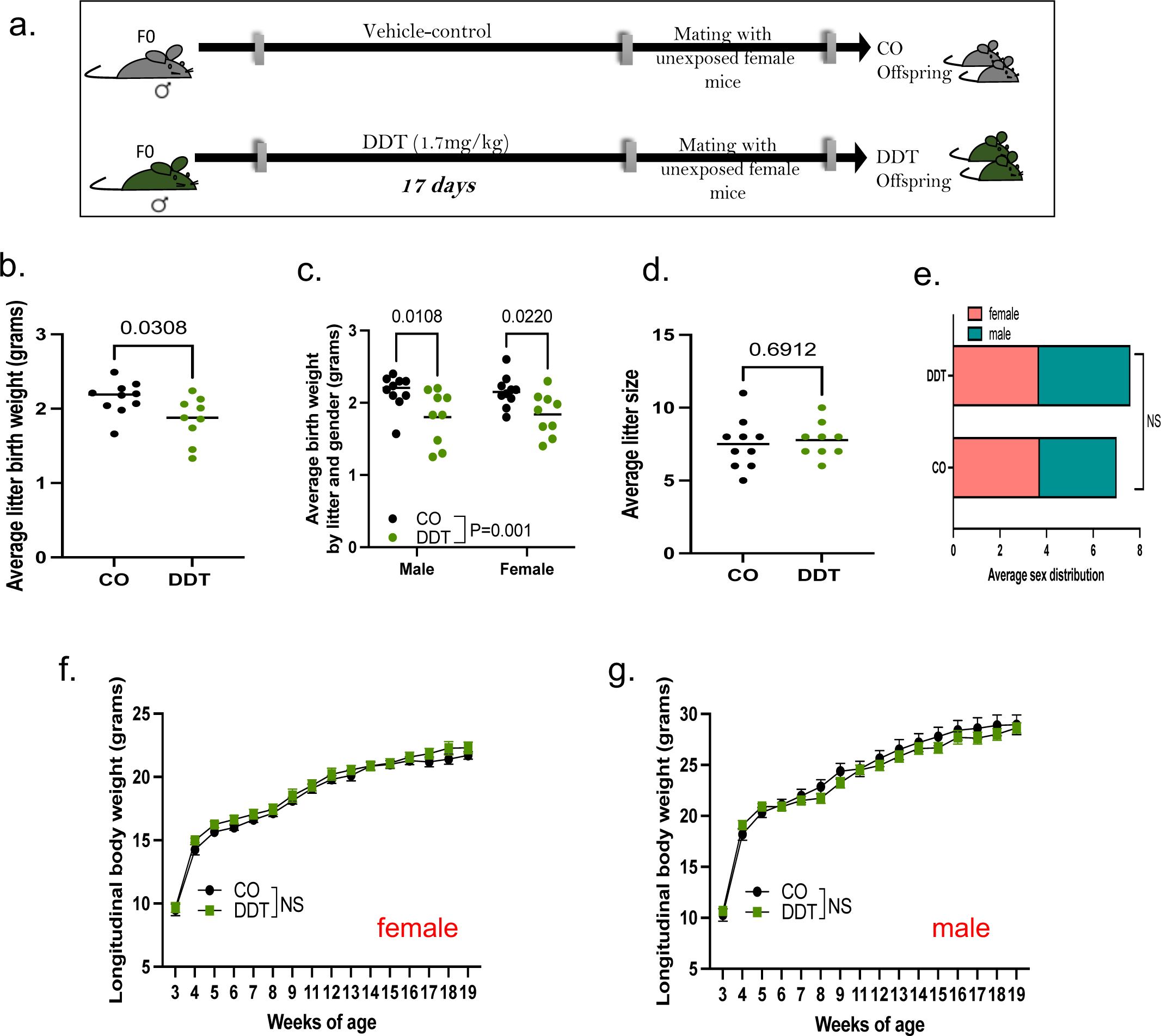
Pre-conception paternal DDT programs offspring’s birth weight. (a) Schematic of experimental design: Control (CO) and DDT-exposed male mice were mated to unexposed females to produce offspring. (b) Birth weight (grams) of CO and DDT offspring. (c) Birth weight (grams) segregated by group and sex. (d) Average litter size and (e) sex distribution. (f-g) Longitudinal body weight (3-19 weeks of age) in female (f) and male (g) offspring. Data shown as mean (horizontal bars in scatter plots), n=8-10 litters/group (b-e), n=13-16 males and females/group (f-g). Data was analyzed by t-test (b,d), Fisher’s exact test (e), two-way ANOVA (c) or repeated measures ANOVA (f,g).

### Paternal DDT exposure impairs placenta growth and development

Given the small birth weight in DDT offspring, we next examined the potential impact of pre-conception paternal DDT exposure on fetal growth and placenta development. Pregnant mouse dams mated with either DDT-exposed or control males were euthanized and both placenta and fetal tissues harvested on embryonic day(E) 13.5, after placentation is complete in mice (**Fig. 2a**). In agreement with the low birth weight, fetal weights were also reduced with a significant decrease in male fetuses only (**Fig. 2b-c)**. We also detected a decrease in placenta weights, again with more pronounced effects in the male placentas (**Fig. 2d-e**).

**Figure 2.**
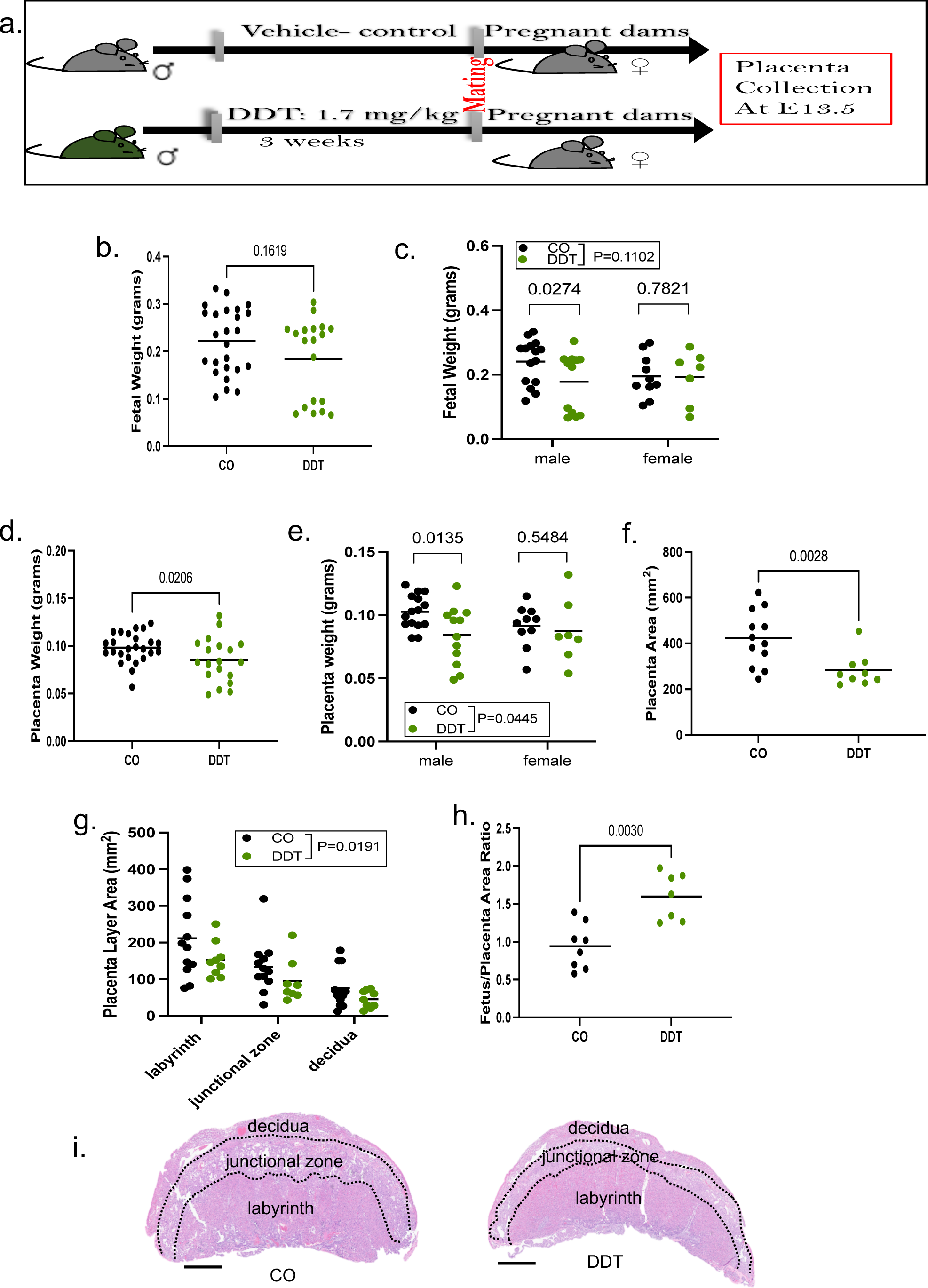
Paternal DDT exposure impairs fetal and placental growth. (a) Schematic of experimental design: Control (CO) and DDT-exposed male mice were mated to unexposed females to produce offspring. Pregnant dams were euthanized on E13.5 for placenta collection. (b-c) Fetal weight (grams) by (b) group and by (c) group and sex. (d-e) Placenta weight (grams) by (d) group and by (e) group and sex. (f) Placental total area, (g) placenta layer-specific area and (h) fetus/placenta ratio. (i) Representative picture of mouse H&E-stained placenta showing layers; bar=0.8 mm. Images were analyzed via image J software. Data shown as mean (horizontal bars in scatter plots), n=19-25/group (b-e), n=9-12/group (f-g), n=8-7/group (h). Data was analyzed by t-test (b, d, f, h) or two-way ANOVA (c, e, g).

Next, we examined placenta morphology and area using H&E-stained sections. In line with the smaller mass observed in placentas of DDT offspring, their placentas also had decreased total area with a higher fetus to placenta area ratio compared to controls (**Fig. 2f-h**). It has been reported that defects in both the junctional zone (JZ) and the labyrinth zone are associated with fetal growth restriction^2^. Consistent with that, we also detected a layer-specific reduction in placentas of DDT offspring (**Fig. 2g**).

### Paternal DDT exposure reduces placenta glycogen content and expression and activity of O-GlcNAc transferase (OGT)

Glucose is the major nutrient source for energy generation in the placenta. The glycogen cells within the placenta provides energetic and hormonal support to the fetus^2^. In agreement with the smaller size of DDT placentas, we found a significant reduction in placental glycogen levels assessed via PAS/amylase assay (**Fig. 3a-c**). This phenotype was again sex-specific with a more pronounced decrease in male placentas from the DDT group (**Fig. 3b**).

**Figure 3.**
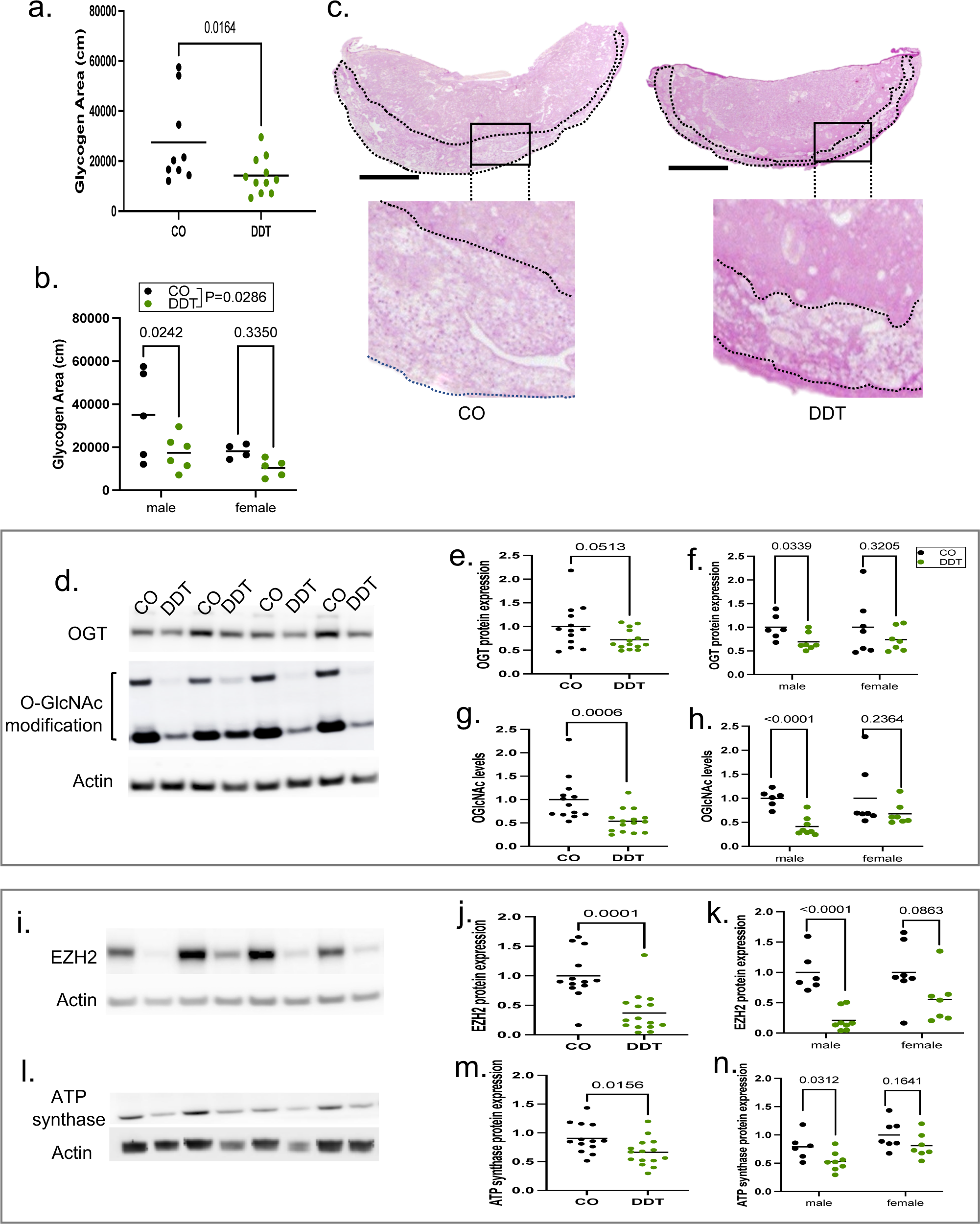
Paternal DDT exposure reduces placenta expression and activity of O-GlcNAc transferase (OGT). Pregnant dams were euthanized on E13.5 for placenta collection. (a-b) Placenta glycogen levels (via PAS-amylase staining) by (a) group and by (b) group and sex. (c) Representative picture of mouse PAS-amylase stained mouse placentas showing area containing glycogen trophoblast cells (delineated by dashed lines). Inset showing glycogens storing cells (glycogen stores appear white in section); bar=1mm; Images were analyzed via image J software. (d-h) Placenta OGT expression and O-GlcNAc levels: (d) Representative western-blots showing OGT expression and O-GlcNAcylation levels with beta-actin as loading control; (e-h) Quantification of placenta OGT and O-GlcNAc levels (e,g) by group and (f,h) by group and sex. (i-k) Placenta EZH2 expression levels: (i) Representative western-blots showing EZH2 expression with beta-actin as loading control; (j-k) Quantification of placenta EZH2 expression (j) by group and (k) by group and sex. (l-n) Placenta ATP synthase expression levels: (l) Representative western-blots showing ATP synthase expression with beta-actin as loading control; (m-n) Quantification of placenta ATP synthase expression (m) by group and (n) by group and sex. Data shown as mean (horizontal bars in scatter plots), n=9-10/group (glycogen levels); n=13-15/group (Western-blots). Data was analyzed by t-test (a, e-h, j-k, m-n) or two-way ANOVA (b).

Reasoning that reduced glycogen levels could disrupt energy sensor activity in placenta, we next examined the levels and activity of OGT (O-Linked N-Acetylglucosamine [GlcNAc] transferase). This enzyme is a nutrient sensor and its activity is dependent on glucose flux through the hexosamine biosynthetic pathway^17 18^. OGT plays an important role in placenta development and function ^19^. Indeed, we found that both OGT protein levels and OGT-catalyzed protein O-GlcNAcylation were reduced in placentas of DDT offspring (**Fig.3d-h**).

OGT is an X-linked gene. Because X chromosome inactivation is malleable in the placenta^20^, and this tissue displays sexual dimorphism in X-linked genes, we next examined whether there were differences in OGT expression and function by gender. Indeed, we found that paternal DDT associated reduction in placental OGT expression and O-GlcNAcylation was sex-specific and more pronounced in male placentas (**Fig.3f-h**). Importantly, sexual dimorphism in the expression of OGT has also been reported in human placentas^6^.

As a O-GlcNAc transferase, OGT plays a key role in many biological processes via posttranslational modification and stabilization of both cytosolic and nuclear proteins^18^. For instance, OGT controls chromatin remodeling by mediating histone methylation via stabilization of the histone methyltransferase EZH2^21^. Consistent with lower levels of OGT and O-GlcNAcylation, levels of EZH2 protein were significantly reduced (**Fig.3i-k**) in placentas of DDT offspring, particularly in male placentas.

Another documented target of OGT is ATP synthase^22^, a protein that controls energy levels in the form of ATP to fulfill the bioenergetic function of the placenta. Although not as pronounced as EZH2, we also found a sex-specific reduction of ATP synthase protein levels, with a more marked decreased in DDT male offspring placentas compared to controls (**Fig.3l-n**).

### Paternal DDT exposure programs metabolic dysfunction in offspring

Small birth weight has been associated with chronic diseases such diabetes and other diseases later in life^23^. Given this relationship, we next examined whether paternal DDT exposure increased metabolic dysfunction in the progeny ages 8-10 week of age. Indeed, DDT offspring showed poor metabolic function in adulthood, with a sex-specific effect (**Fig. 4a-e**). Male offspring displayed severe glucose intolerance and a subtle, but significant, increase in insulin resistance compared to controls (**Fig. 4b,d**). However, no differences in metabolic function were observed between DDT and CO female offspring of the same age (**Fig. 4c,e**).

**Figure 4.**
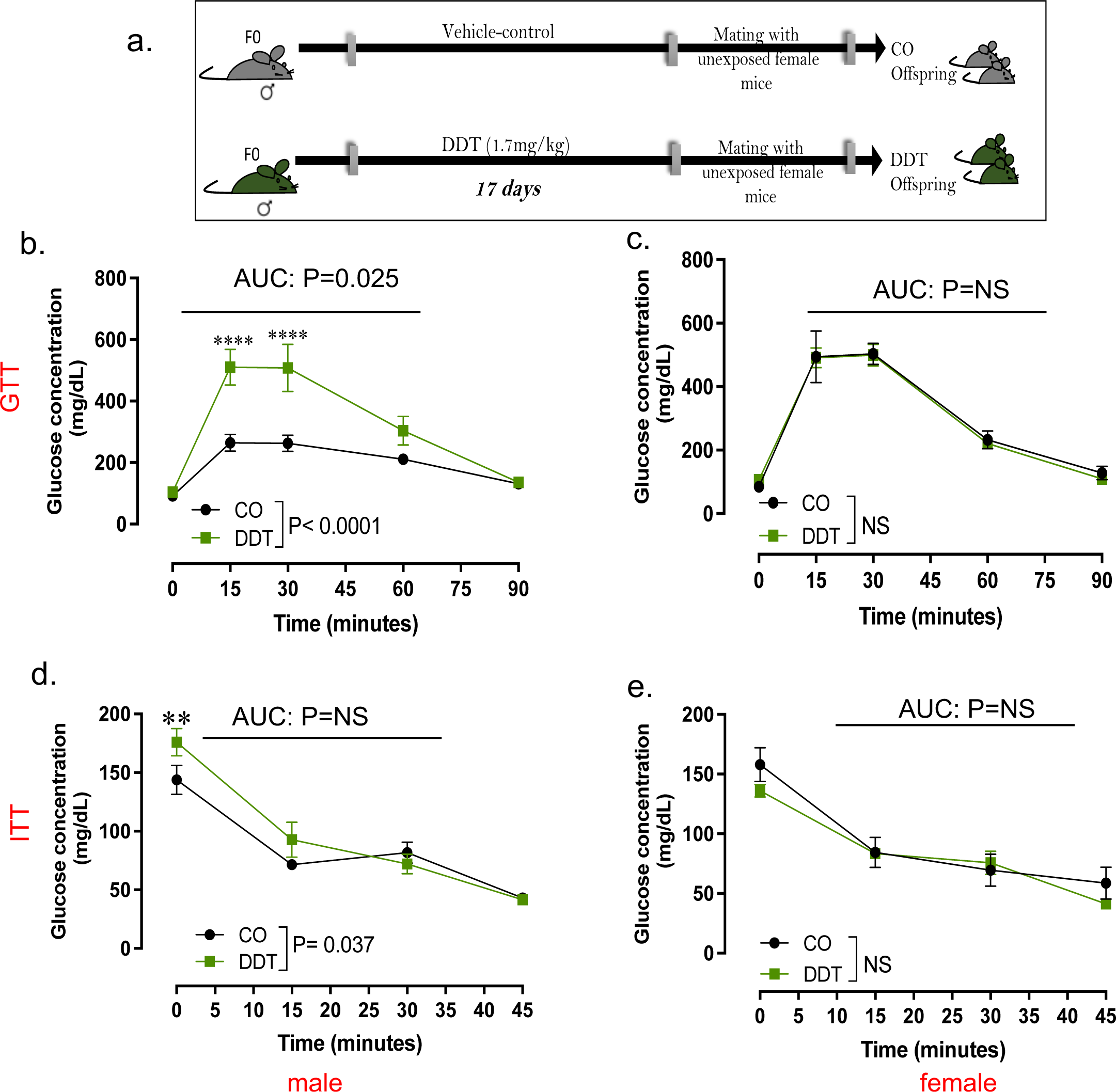
Paternal DDT exposure programs metabolic dysfunction in offspring. (a) Schematic of experimental design: Control (CO) and DDT-exposed males were mated to unexposed females to produce offspring. Metabolic function in offspring of CO and DDT-exposed males at 8-10 weeks of age: (b-c) Glucose tolerance test (GTT) and (d-e) Insulin Tolerance Test (ITT) in male (b,d) and female (c,e) offspring. Data shown as mean±SEM, n=9 males and females /group (b,c), n=8-9 males and females /group (d,e). Data was analyzed by repeated measures ANOVA (b-e). AUC data was analyzed by t-test (b-e).

### Placenta-specific genetic deletion of OGT partially replicates the effects of paternal DDT on offspring metabolic function in a sex specific manner

To examine whether suppression of OGT activity in placenta was functionally linked to programming of metabolic dysfunction in DDT progeny, we used a mouse Cre -Lox system for placenta-specific genetic deletion of OGT. Female offspring with placenta-specific OGT reduction (OGTKO or OGThet) showed no signs of metabolic dysfunction compared to control littermates, in agreement with our finding in DDT female offspring (**Fig. 5a-b**). Though male offspring with placenta-specific OGT deletion (OGTKO/Y) genotype showed metabolic dysfunction, their phenotype only partially replicated that of DDT male offspring. While placenta OGTKO/Y male offspring showed a subtle increase in insulin resistant in early adulthood (around 12 weeks of age), they showed better glucose handling than WT control male littermates (**Fig. 5c-d**).

**Figure 5.**
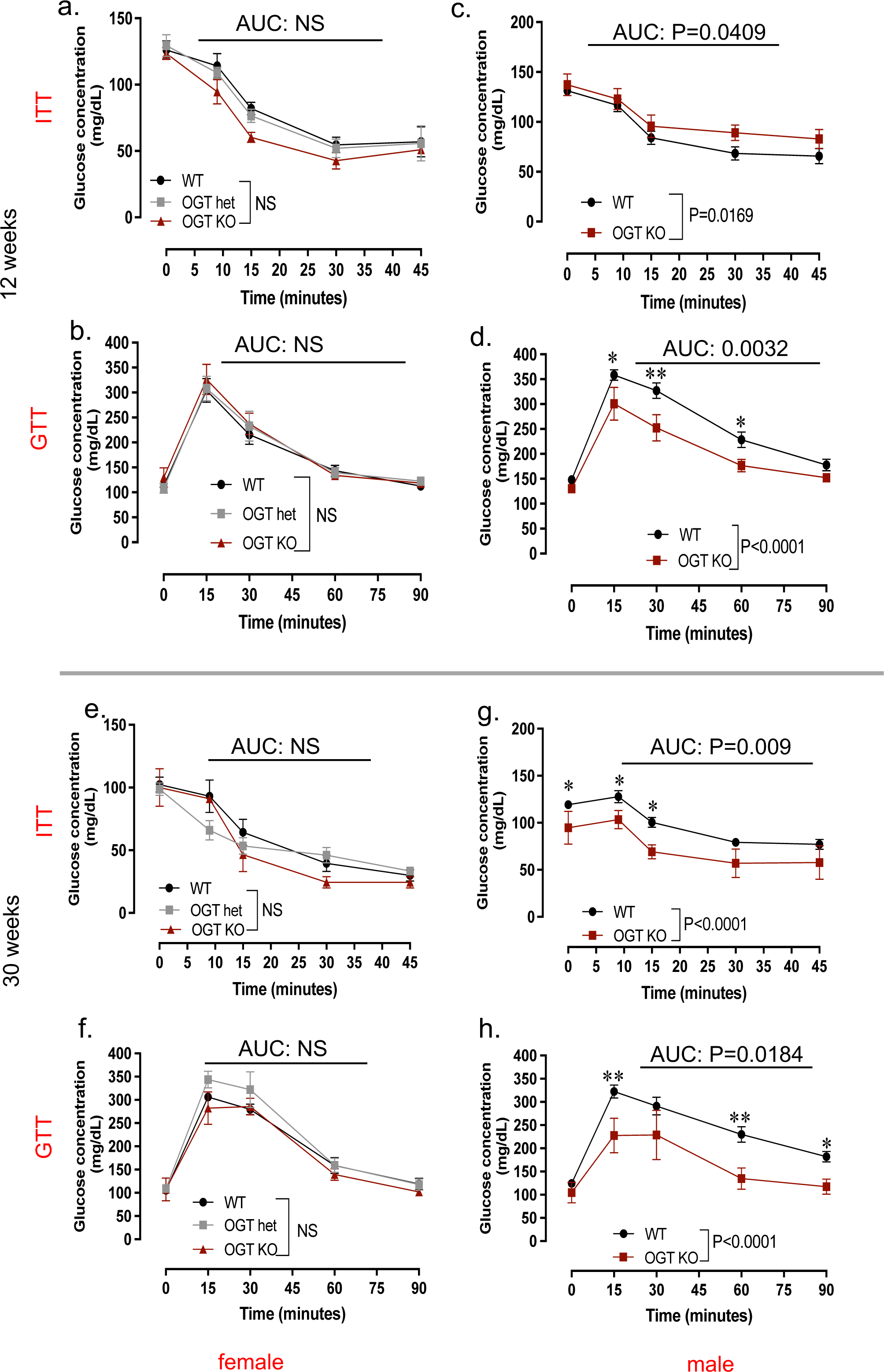
Placenta-specific genetic deletion of OGT does not fully replicate the effects of paternal DDT on offspring metabolic function. (a-d) Metabolic function in WT and OGTKO mice at 12 weeks of age: (a,c) Insulin tolerance test (ITT) and (b,d). Glucose Tolerance Test (GTT) in female (a,b) and male (c,d) offspring. (e-h) Metabolic function in WT and OGTKO mice at 30 weeks of age: (e,g) Insulin tolerance test (ITT) and (f,h) Glucose Tolerance Test (GTT) in female (e,f) and male (g,h) offspring. Data shown as mean ± SEM in all graphs; (a,c) n=21 (WT males and females), n=9 (OGTKO males and females), n=5 (OGThet); (b,d) n=19 (WT males and females), n=9 (OGTKO males and females), n=6 (OGThet females); (e-h) n=16 (WT males and females), n=5 (OGTKO males and females), n=8 (OGThet females); ITT and GTT data was analyzed by repeated measures ANOVA (a-h). AUC data was analyzed by t-test (a,b,e,f) or one-way ANOVA (c,d,g,h).

Because metabolic dysfunction tends to worsen with age^24^, we repeated metabolic assessments at 30 weeks of age. Metabolic function in placenta OGT KO or OGThet female offspring did not differ from WT littermates (**Fig. 5e-f**). However, placenta OGTKO/Y males showed better glucose tolerance and insulin sensitivity compared to WT males (**Fig. 5g-h**).

### Placenta-specific genetic deletion of OGT impacts offspring body-weight gain with age in a sex specific manner

To better understand what metabolic phenotypes observed in offspring with placenta-specific OGT reduction, we evaluated their body weight compared to WT mice from weaning to 30 weeks of age (**Fig. 6a-d**). Not surprisingly, there were no differences in body weight in placenta OGTKO or OGThet female offspring compared to WT controls (**Fig. 6a**). However, though placenta OGTKO/Y and WT males displayed similar body weight gain until about 12 weeks of age, OGTKO/Y males did not gain as much body weight after this time point compared to WT controls, weighing significantly less by 24 weeks of age (**Fig. 6b**).

**Figure 6.**
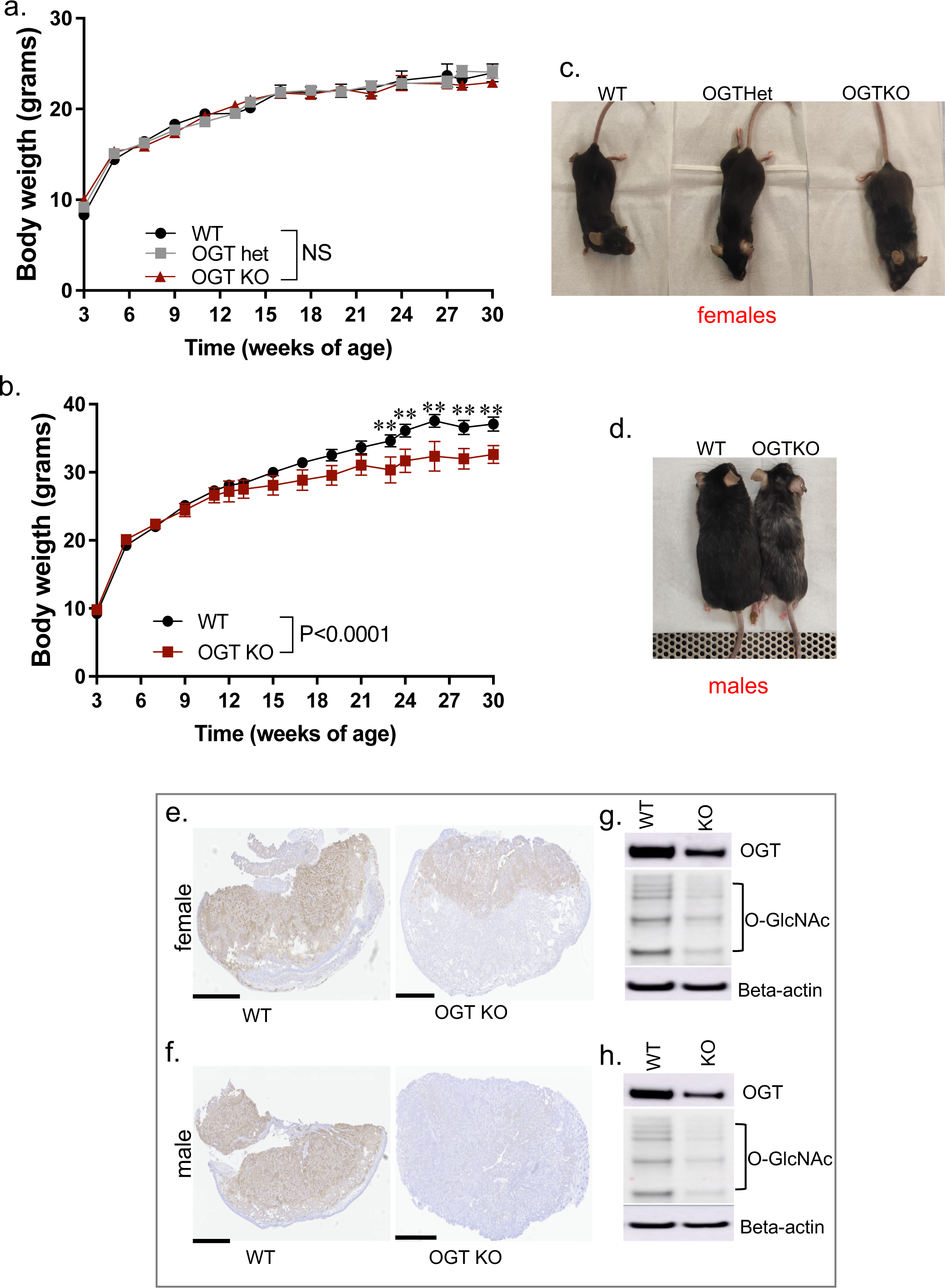
Placenta-specific genetic deletion of OGT impacts offspring’s bodyweight in a sex specific manner. (a-b) Longitudinal body weight (3-30 weeks of age) in (a) WT, OGThet or OGTKO females and in (b) WT or OGTKO males. Representative pictures showing (c) WT, OGThet or OGTKO females and (d) WT or OGTKO males at 30 weeks of age. (e-f) Representative pictures showing OGT protein immunostaining in placentas of (e) females or (f) males with a WT or OGTKO genotype; bar=1mm. (g-h) Representative western-blots showing OGT protein expression and O-GlcNAcylation levels in placentas of (g) females or (h) males with a WT or OGTKO genotype; beta-actin, loading control. Data shown as mean *±* SEM(a-b); n=16 (WT males and females), n=5 (OGTKO males and females), n=8 (OGThet females). Body weight data was analyzed by repeated measures ANOVA.

Because OGT is an X-linked gene, we wondered whether placenta Cre recombinase mediated OGT deletion is more efficient in males than in females in this model. We found that this is indeed the case. While placenta OGTKO female showed decreased OGT expression and activity (O-GlcNAcylation levels) compared to the WT group (**Fig. 6e-g**), they still retained higher expression levels of OGT protein compared to OGTKO/Y male placentas, where OGT protein expression and activity is almost completely lost (**Fig. 6f-h**).

## Discussion

Using a mouse model, we found that preconception paternal DDT exposure is linked to a decrease in the progeny’s birth weight and reduced placental size. We also observed a placenta-specific reduction of glycogen levels and the nutrient sensor and epigenetic regulator OGT, with more pronounced phenotypes observed in male placentas. The progeny of DDT-exposed males also exhibited sexual dimorphism in metabolic function, with DDT male offspring developing both glucose intolerance and insulin resistance. However, placenta-specific genetic reduction of OGT only partially replicates the metabolic phenotype observed in offspring of DDT-exposed males.

Our findings are consistent with epidemiological studies linking parental exposure to persistence organic pollutants and other environmental chemicals to low birth size in offspring in humans^13-16,25^. In our model, the reduced birth weight in offspring of DDT-exposed males was associated with impaired fetal growth and placenta size.

There is accumulating evidence for placenta as a mediator of environmentally-induced intergenerational transmission of disease^3^. Given the tight connection between the fetus and the mother during gestation, it is well documented that maternal experiences in pregnancy impact placenta development and function ^5,6^. In our paradigm, paternal exposure to the environmental toxicant DDT also impaired placenta development, its glycogen storage and nutrient sensing signaling.

In humans and mice, the inner cell mass and the trophectoderm— the layer containing cells that give rise the placenta and other extraembryonic tissues— are the first definitive embryonic cell lineages to appear^3^. The integrity and function of the trophectoderm is critical for the blastocyst-stage embryo implantation and pregnancy success. Defective placentation can result in fetal growth restriction and other pregnancy complications^26,27^. Although details need to be elucidated, our data suggest that pre-conception paternal environmental exposures can contribute to disruption of this process and lead to defective placentation.

Abnormal placenta function and adverse events in fetal development have been associated with disease later in life. In this study, we found a paternal exposure to DDT also results in metabolic dysfunction and poor glucose handling in the progeny. This effect is sex specific and the phenotype more severe in male offspring. Interestingly, both human and animal studies link maternal exposure to DDT to metabolic dysfunction in the next generations^28-30^. These findings, along with previously published reports, suggest that paternal exposures may be as important as maternal ones regarding determination of the progeny’s phenotypes. This striking overlap between the impact of maternal or paternal exposure on offspring phenotypes has been noted by others ^31^, and our current findings raise the possibility that the placenta may be the common mediator of intergenerational epigenetic inheritance of phenotypes.

OGT is encoded by an X linked-gene and is a key regulator of several biological processes through its posttranslational modification of both nuclear and cytosolic proteins. Unlike other somatic tissues, the inactivation of the X chromosome in the placenta is plastic, allowing for its reactivation in response to stress^20,32^. It has been proposed that this is the reason why female fetuses are more resilient to environmental stressors^33^. Our findings are in line with this concept as paternal DDT exposure has greater impact on placenta of male fetuses and suppresses placenta OGT signaling in a sex-specific manner. Importantly, this sexual dimorphism in placenta OGT expression has also been reported for maternal exposures^6^, further strengthening the notion that placenta abnormalities could mediate both paternal and maternal effects on the progeny.

In our study, placenta OGT reduction did not fully replicate the metabolic dysfunction observed in DDT male offspring. It is possible that the divergent metabolic phenotypes between DDT male offspring and placenta OGT KO offspring is due to timing. While in the placenta OGTKO mouse model use here, Cre-induced genetic knockdown occurs early in development^34^, it’s possible that OGT suppression in placenta of DDT offspring is a later event in the placentation process, though this still needs to be investigated.

OGT and its biochemical mark, O-GlcNAcylation, play a role in placentation through its regulation of syncytiotrophoblast formation. In BeWo trophoblast cells, OGT depletion via siRNA leads to spontaneous differentiation^35^. Although it does not completely phenocopy the poor glucose handling in DDT male offspring, it is possible that OGT genetic reduction in the early trophectoderm causes a more serious disruption in placentation than observed in the paternal DDT paradigm. This likely leads to suboptimal fetal development and more severe health phenotypes such as impaired body weight gain and neurodevelopmental disorders^6^ later in life. While not described in this report, our early findings suggest that placenta OGT genetic reduction results in in many other health alterations in offspring as they age.

Why OGT genetic reduction results in sex specific effects on metabolic homeostasis is not entirely clear. However, the fact longitudinal body weight of OGTKO and OGThet females does not differ from the control littermates unlike OGTKO/Y males offers a clue. We also showed that OGTKO females retain some degree of OGT expression in placenta, which could be enough to provide protection. It has been reported that placenta EZH2, an OGT target, is responsible for the female fetus resilience to maternal stress ^36^. However, whether this epigenetic modifier plays a role in our model still need to be investigated.

Employing the same placenta-specific OGTKO mouse model used here, a recent study^37^ found no differences in glucose homeostasis in males 6-8 weeks of age or fed a high-fat diet, but reported an increase in insulin sensitivity in OGThet placenta females fed a 12-week course of high fat diet. This was associated with increased hepatic insulin/Akt axis signaling. The study also reported a decrease in pancreatic islet size for mice with placenta-specific OGT knockout compared to the controls but this was compensated for by an increase in the total number of islets. While both of our studies found better glucose handling in mice with placenta-specific OGT reduction, this phenotype was detected exclusively in males in the present study. Differences between our studies likely stem from the timing of metabolic testing and different dietary conditions. In the present study, mice were kept on chow diet throughout the study while in the Moore et al study, mice were switched to a high-fat diet at 12 weeks of age. We also backcrossed the original FBV Cyp19-Cre mice into the c57bl6 background, which could further explain the phenotypic differences between the two studies.

Finally, while we showed a link between paternal DDT exposure and programming of placenta development, the environmentally-induced paternal factors involved in placenta development need to be fully evaluated. In our model, mating of DDT-exposed males with unexposed females was performed overnight and offspring and placenta traits likely stem from changes in the sperm epigenome as reported before. The impact of environmental factors on the sperm epigenome, particularly sperm RNAs, has been well documented by us and others^38-42^. However, whether and how the sperm epigenome plays a role in placenta development remains to be investigated.

Despite many remaining questions, our study show that preconception paternal DDT induced alteration in offspring is linked to placenta defects. Our findings, along with recent reports^7,9^, raise the tantalizing possibility that placenta programming is a mediator of environmentally induced intergenerational epigenetic inheritance of phenotypes and needs to be further evaluated.

## Materials and Methods

### DDT exposure and breeding

The C57BL/6 strain of mice (Taconic Biosciences) was used in this experiment. Adult male mice (8 weeks of age) were either treated daily with a DDT solution (1.7 mg/kg) or vehicle-control solution (peanut oil) via oral gavage for 17 days. Duration of exposure was chosen in order to encompass the entire period of post-testicular sperm transit in the epididymis (which takes about 15 days in mammals), the last stage of sperm maturation ^43^. The DDT solution used mimics the formulation of DDT before its ban in the U.S.: 77.2% p,p’-DDT and 22.8% o,p’-DDT as described before ^29^.

At the end of the exposure period, DDT or vehicle-control exposed male mice were mated overnight to unexposed females to generate the offspring. Pregnant dams were allowed to deliver their offspring or euthanized on embryonic day (E)13.5 for placenta collection as described below. All mice utilized in this study were kept on a standard chow diet including the exposures and breeding periods, for the extent of pregnancy (21 days) and after birth. To avoid litter-effect, pups were cross-fostered one day after birth. All pups were weaned on postnatal day (PND) 21. The offspring of control-vehicle or DDT-exposed fathers were used to study metabolic function as described in the following sections.

The number of offspring/litters stemming from different fathers included in each experimental endpoint is shown is **Table S1.**

All animal procedures were approved by the Georgetown University Animal Care and Use Committee, and the experiments were performed following the National Institutes of Health guidelines for the proper and humane use of animals in biomedical research. Animals were randomized to each exposure group and treatments were performed blindly.

### Placenta harvest

Non-exposed female mice were mated with DDT or vehicle-control exposed males overnight as described above. Mating was confirmed by detection of a vaginal plug. This was considered the first day of pregnancy or embryonic day (E) 0.5. Pregnant dams were housed in groups of two with free access to food and water and euthanized either on **E13.5**, after placentation is complete in mice. Placentas and fetuses were weighed upon dissection and either snap frozen or fixed in 10% neutral-buffered formalin. DNA from fetal tail tips were used for gender determination via commercial genotyping (Transnetyx, Inc.) of the Y chromosome specific gene, SRY.

### Placenta morphometric, cellular and molecular assessments

Placenta area and structures were evaluated on H&E-stained sections collected at using the Image J software. Morphometric ends points were total placental area and area of each placental layer (labyrinth, junctional zone and maternal decidua). All comparisons were made using mid-sagittal sections.

Glycogen cells within the junctional zone were stained with periodic acid–Schiff (PAS), followed by digestion without and with α-amylase, using a commercial PAS staining system (Sigma-Aldrich, catalog #395), according to manufacturer’s instructions. Protein levels of OGT, EZH2, ATP and patterns of protein O-GlcNAcylation were assessed via western-blot (as described below) in placenta (See **Table S2** for list antibodies and dilutions).

The number of samples/litters stemming from different fathers included in placenta endpoint is shown is **Table S1.**

### Assessment of longitudinal body weight and metabolic parameters in offspring

Offspring of vehicle-control or DDT exposed males were weighed one day after birth and weekly starting at weaning (3 weeks of age) and monitored until 20 weeks of age.

Glucose tolerance test (GTT) and insulin tolerance test (ITT) were performed in 8 to10 week-old mice. For the GTT, mice were fasted for 6 hours, injected i.p. with glucose (2.5g/kg) and glucose measurements taken at 0, 15, 30, 60, and 90 min using a Accu-check meter. For the ITT, mice were fasted for 4 hours, injected i.p. insulin (0.75 U/kg), glucose measurements at 0, 15, 30, 45, 60 and 90 min using an Accu-check meter.

The number of offspring/litters stemming from different fathers included in metabolic endpoints is shown is **Table S1.**

### Placenta-specific genetic reduction of OGT

A mouse Cre-Lox system under the control of the Cyp19-Cre promoter was used in this experiment. For the placenta-specific reduction of OGT, Ogt^tm1Gwh^/(X^OGTflox^/X^WT^); CYP19-Cre (placental-Cre recombinase homozygote, P. Cre^+/+^)] females were mated to hemizygous [Ogt^tm1Gwh^/(X^OGTflox^/Y)] males, resulting in offspring of the following genotypes as described before^44^: **functional WT** females (X^WT^/X^WT^/Cre^+^ or X^WT^/X^WT^/Cre^−^ or X^OGT^/X^WT^/Cre), **OGThet** female (X^OGT^/X^WT^/Cre^+^), **OGTKO** female (X^OGT^/X^OGT^/Cre^+^), functional **WT/Y** males (X^WT^/Y/Cre^+^ or X^WT^/Y/Cre^−^) or male **OGTKO/Y** (X^OGT^/Y/Cre^+^). In this mouse model, CYP19-Cre is expressed throughout the junctional and labyrinth trophoblast in placentas^34^. CYP19-Cre mice (*FVB* strain) used in this study was a gift from Dr. Gustavo Leone from the Medical College of Wisconsin. These mice were crossed back to a C57BL/6 strain for at least 5 generations before being use in this study. The Ogt^tm1Gwh^ mice were obtained from Jackson laboratories (Strain #:004860) and have a B6.129 (C57BL/6; 129P3/J) mixed background.

Resulting offspring were used for body weight and metabolic studies as described in the next section. Another cohort of pregnant dams were euthanized before giving birth and used for placenta collection to confirm OGT genetic reduction via western blot or immunohistochemistry (described in the next sections). Offspring’s genotyping (OGT and SRY) was carried out commercially (via tail clipping) by Transnetyx, Inc.

### Longitudinal assessment of body weight and metabolic parameters in placenta specific WT or OGTKO offspring

Offspring were weighted weekly starting at weaning. Glucose tolerance test (GTT) and insulin tolerance test (ITT) were performed at 11-12 weeks of age and then again at 30 weeks of age. For the GTT, mice were fasted for 6 hours, injected i.p. with glucose (2.5g/kg) and glucose measurements taken at 0, 15, 30, 60, and 90 min using a Accu-check meter. For the ITT, mice were fasted for 4 hours, injected i.p. insulin (0.75 U/kg), glucose measurements at 0, 9, 15, 30 and 45 min using an Accu-check meter.

### Western-blots

Total protein was extracted from mammary tissues using RIPA buffer with Halt™ Protease Inhibitor Cocktail (Thermo Fisher). Protein extracts were resolved on a 4-12% denaturing polyacrylamide gel (SDS-PAGE). Proteins were transferred using the iBlot® 7-Minute Blotting System (Invitrogen, USA) and blocked with 5% non-fat dry milk for 1h at room temperature. Membranes were incubated with the specific primary antibodies (for antibody specifications and dilutions see **Table S2**) at 4°C overnight. After several washes, the membranes were incubated with horseradish peroxidase (HRP)-conjugated secondary antibody at room temperature for 1h. Membranes were developed using the Chemiluminescent HRP antibody detection reagent (Denville Scientific Inc., USA), and exposed to blot imaging systems (Amersham^TM^ Imager 600, GE Healthcare Life Sciences). Optical density of the bands was quantified using Quantity-one software (BIO-RAD, USA). To control for equal protein loading, expression of the proteins of interest was normalized to the β-actin signal.

### Immunohistochemistry

Five microns sections from formalin fixed paraffin-embedded placenta tissues were de-paraffinized with xylenes and rehydrated through a graded alcohol series. Heat induced epitope retrieval (HIER) was performed by immersing the tissue sections in Target Retrieval Solution, Low pH (DAKO) in the PT Link (DAKO). Immunohistochemical staining was performed using a horseradish peroxidase labeled polymer (Agilent, K4003) according to manufacturer’s instructions. Briefly, slides were either treated with 3% hydrogen peroxide and 10% normal goat serum for 10 minutes each, and exposed to primary antibody for OGT (1:200, Cell Signaling, D1D8Q) for 1hr at room temperature. Slides were exposed to the appropriate HRP labeled polymer for 30min and DAB chromagen (Dako) for 5 minutes. Slides were counterstained with Hematoxylin (Fisher, Harris Modified Hematoxylin), blued in 1% ammonium hydroxide, dehydrated, and mounted with Acrymount. Consecutive sections with the primary antibody omitted were used as negative controls.

### Statistical analysis

Statistical analyses were performed using GraphPad Prism (GraphPad Software, San Diego, CA, USA). Normal probability plots were used to ascertain normality, which were confirmed by Anderson-Darling, D’Agostino-Pearson, Kolmogorov-Smirnov or Shapiro-Wilk tests. Where data failed the normality test, log transformation was performed prior to statistical analysis or a non-parametric test used. Repeated measures two-way ANOVA was used to analyze longitudinal body weight curves, ITT and GTT (group, time). Two-way ANOVA was used to analyze birth weight, placenta weight, glycogens levels (group, sex or layer). Sex distribution was analyzed by Fisher’s exact test. All other end-points were analyzed using unpaired t-test (for two groups) or one-way ANOVA (for three groups). Differences were considered statistically significant at P< 0.05. Unless indicated, *n* corresponds to the number of animals used in each experiment. The number of samples/litters stemming from different fathers included in each endpoint is shown is **Table S1.**

## Acknowledgments

We thank the following Lombardi Cancer Center Shared Resources (SR) for their assistance: Animal Model SR, Histopathology & Tissue SR, Microscopy and Imaging SR. We also thank Ms. M. Idalia Cruz for her technical assistance with animal experiments.

## Author contribution

E.C. and R.S.D.C. supervised the animal work and tissue collection and performed molecular analysis; A.C., M. J., and D. G. P. completed a portion of the animal work and placenta analysis. O.D., G. F. and M.S. assisted with animal studies, processed tissues and/or helped perform molecular analysis; S. P.C. P. contributed with data analysis and wrote the manuscript; S.D.A. conceived the study, oversaw the research and wrote the manuscript.

**The authors declare they have no actual or potential competing financial interests**

## Supplementary Tables

**Table S1.**
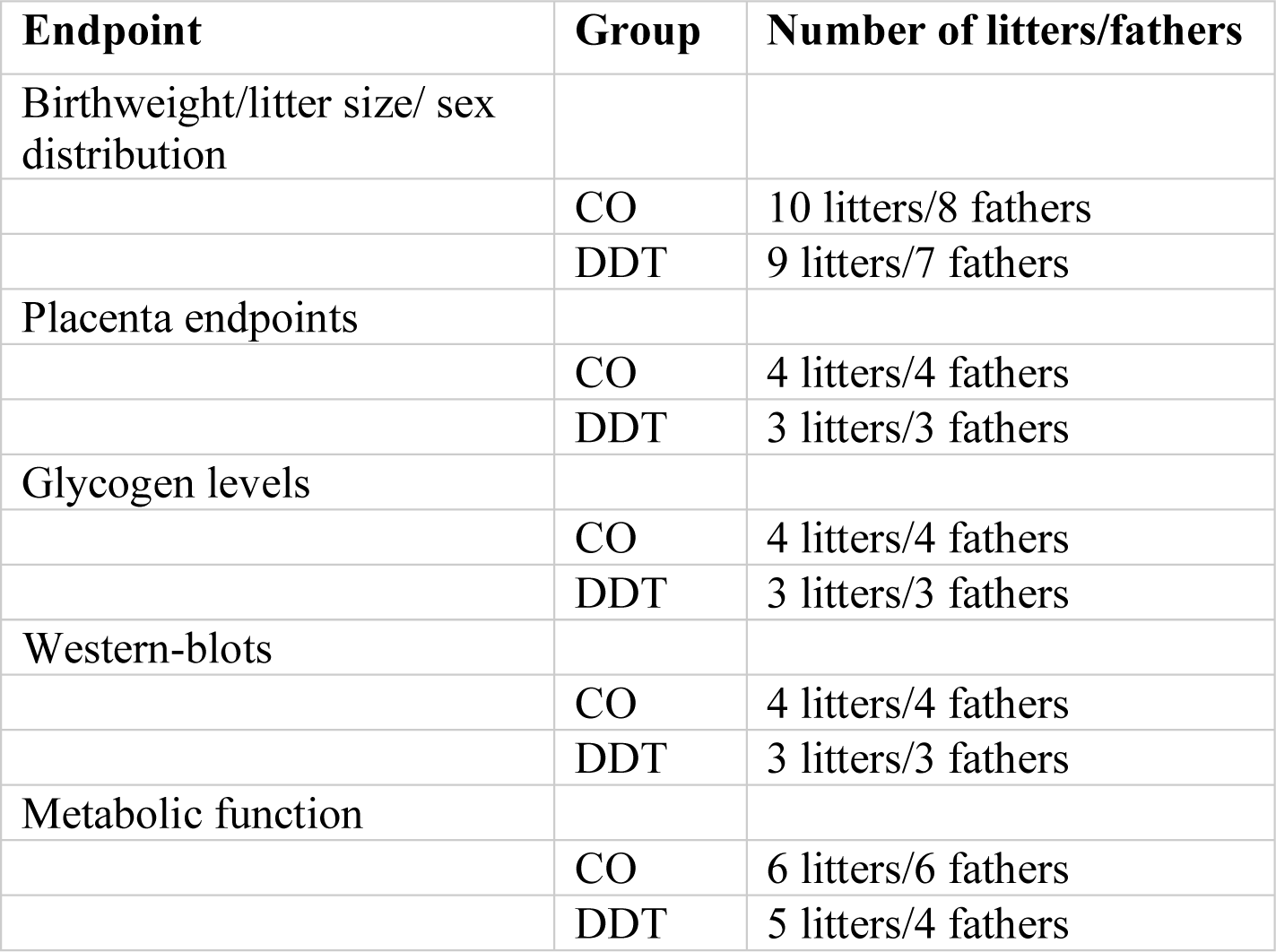
Paternal DDT study: Number of litters and fathers represented in each experimental endpoint.

| Endpoint | Group | Number of litters/fathers |
| --- | --- | --- |
| Birthweight/litter size/ sex distribution |  |  |
|  | CO | 10 litters/8 fathers |
|  | DDT | 9 litters/7 fathers |
| Placenta endpoints |  |  |
|  | CO | 4 litters/4 fathers |
|  | DDT | 3 litters/3 fathers |
| Glycogen levels |  |  |
|  | CO | 4 litters/4 fathers |
|  | DDT | 3 litters/3 fathers |
| Western-blot |  |  |
|  | CO | 4 litters/4 fathers |
|  | DDT | 3 litters/3 fathers |
| Metabolic function |  |  |
|  | CO | 6 litters/6 fathers |
|  | DDT | 5 litters/4 fathers |

**Table S2.** List of Antibodies Used for Western-blots or Immmohistochemistry (IHC)

| Company | Catalog # | Antibody | Application/Dilution |
| --- | --- | --- | --- |
| Abcam | ab177941 | Anti-OGT / O-Linked N-Acetylglucosamine Transferase Monoclonal Antibody (EPR12713) | Wester-blot<br>1:1000 |
| Cell Signaling Technology | 24083 | Anti-OGT/ O-Linked N-Acetylglucosamine Transferase Monoclonal Antibody(D1D8Q) | IHC<br>1:200 |
| Santa Cruz Biotechnology | SC-59623 | Anti-O-GlcNAc Monoclonal Antibody (CTD110.6) | Wester-blot<br>1:1000 |
| Cell Signaling Technology | 5246 | Anti-EZH2 Monoclonal Antibody (D2C9) | Wester-blot<br>1:1000 |
| ProteinTech | 66037 | Anti-ATP5A1 Monoclonal Antibody | Wester-blot<br>1:5000 |
| Abcam | ab8229 | Anti-Beta Actin Polyclonal Antibody | Wester-blot<br>1:1000 |

